# Assessment of the Glycan-Binding Profile of *Pseudomonas aeruginosa* PAO1

**DOI:** 10.1101/2023.04.20.537720

**Authors:** Hector Sanchez, George A. O’Toole, Brent Berwin

## Abstract

*Pseudomonas aeruginosa* is an opportunistic pathogen that can establish acute and chronic infections in individuals that lack fully functional innate immunity. In particular, phagocytosis by neutrophils and macrophages is a key mechanism that modulates host control and clearance of *P. aeruginosa*. Individuals with neutropenia or cystic fibrosis are highly susceptible to *P. aeruginosa* infection thus underscoring the importance of the host innate immune response. Cell-to-cell contact between host innate immune cells and the pathogen, a first step in phagocytic uptake, is facilitated by simple and complex glycan structures present at the host cell surface. We have previously shown that endogenous polyanionic N-linked glycans localized to the cell surface of phagocytes mediate binding and subsequent phagocytosis of *P. aeruginosa*. However, the suite of glycans that *P. aeruginosa* binds to on host phagocytic cells remains poorly characterized. Here we demonstrate, with the use of exogenous N-linked glycans and a glycan array, that *P. aeruginosa* PAO1 preferentially attaches to a subset of glycans, including a bias towards monosaccharide versus more complex glycan structures. Consistent with these findings, we were able to competitively inhibit bacterial adherence and uptake by the addition of exogenous N-linked mono- and di-saccharide glycans. We discuss of findings in the context of previous reports of *P. aeruginosa* glycan binding.

**IMPORTANCE:** *P. aeruginosa* binds to a variety of glycans as part of its interaction with host cells, and a number of *P. aeruginosa-*encoded receptors and target ligands have been described that allow this microbe to bind to such glycans. Here we extend this work by studying the glycans used by *P. aeruginosa* PAO1 to bind to phagocytic cells and by using a glycan array to characterize the suite of such molecules that could facilitate host cell-binding by this microbe. This study provides an increased understanding of the glycans bound by *P. aeruginosa*, and furthermore, provides a useful dataset for future studies of *P. aeruginosa-*glycan interactions.

## INTRODUCTION

*Pseudomonas aeruginosa* is a Gram-negative, opportunistic bacterium that is responsible for a variety of human infections, particularly in individuals from immunocompromised communities. *P. aeruginosa* is capable of losing flagellar swimming motility during chronic infection, likely secondary to forming a biofilm, which in turn allows this microbe to evade immune responses via phagocytic resistance and antibiotic tolerance (1–5). Persons with cystic fibrosis (CF) and neutropenia (lacking neutrophils) are highly susceptible to bacterial infections due defects in phagocytic function necessary for clearance of *P. aeruginosa* (6, 7).

Phagocytosis by neutrophils and macrophages provides the host with an effective defense mechanism against bacterial infection (8). Contact between the host immune cell and the microbe is required for initiation of phagocytosis to proceed. We previously identified that exogenous treatment of phagocytes with a negatively charged phosphoinositide, PIP_3_, promotes phagocytosis of nonmotile *P. aeruginosa* by increasing binding to phagocytes (9). These studies, which suggested that *P. aeruginosa* binds to clustered polyanions, guided our previous work that revealed that endogenous negatively charged glycosaminoglycans (GAGs) can mediate *P. aeruginosa* adhesion. In turn, this observation led to the conclusion that N-linked glycans on phagocytes play a role in bacterial binding and phagocytosis (10). Therefore, a central goal of this study is to further elucidate the initial carbohydrate “handshake” between phagocytic cells and *P. aeruginosa*.

Glycans are found in all of nature and cover cell surfaces by decorating protein and lipid backbones (11, 12). Glycans on host cells are often utilized by pathogenic bacteria for attachment and invasion (13–16), and can even serve as a carbon/energy source (17). The binding of *P. aeruginosa* to glycans has been explored previously in several contexts. We showed that N-linked glycans and glycosaminoglycans on phagocytes can mediate attachment and uptake of *P. aeruginosa* by macrophages (10). N-linked glycans have also been implicated as ligands for bacterial attachment to epithelial cells (18, 19). *P. aeruginosa* can also bind glycan components of mucin (20–23), complex glycans as part of glycolipids (24, 25) and chitin (26, 27).

In this work we further explore the features of N-linked glycan structures that promote *P. aeruginosa* PAO1 binding, including mannose (Man), glucose (Glc), *N*-acetylglucosamine (GlcNAc), galactose (Gal), and fucose. To do so, we utilized single monosaccharide components of an N-linked glycan structure in competition assays. In parallel, we also performed a glycan array study to further evaluate the binding profile of *P. aeruginosa* to different classes of glycans. These studies have led to a better understanding of the glycans that can be bound by *P. aeruginosa* PAO1, and furthermore, provides a useful dataset for others investigating *P. aeruginosa*-glycan interactions.

## RESULTS

### Addition of Exogenous Monosaccharides Compete with Phagocytosis of *P. aeruginosa* by THP-1 Cells

We previously demonstrated that if N-linked glycan synthesis is inhibited on host phagocytic cells the ability of *P. aeruginosa* to bind to the cell surface via these glycans is reduced (10). We hypothesized that the sugars that comprise N-linked glycan structures are necessary for binding and uptake of *P. aeruginosa*. These N-linked glycans structures are primarily composed of mannose, N-acetylglucosamine, fucose, galactose, and sialic acid moieties depending on the cell type, and furthermore, N-linked glycans can be present in different quantities (28). There are three different classes of N-linked glycans (high mannose, complex, and hybrid) and each of these glycans share a common core structure of Man_3_GlcNAc_2_ bound to an asparagine (Asn) residue (29).

To assess binding of glycan structures by *P. aeruginosa* we performed competition assays in which we exogenously supplemented phagocytosis assays with components of N-linked glycans. In this set of experiments, we utilized *P. aeruginosa* PAO1, a motile strain of bacteria previously shown to interact with cell surface glycans (10, 30), and performed gentamicin protection assays with human THP-1 monocytic cell lines, as reported (4, 5, 9, 10), to assess interaction of the microbe with host cells. The assays were performed for 45 minutes, which we have showed previously allows for a reproducible measure of bacterial attachment and subsequent phagocytosis, with minimal change in viability of the internalized bacteria (4, 5, 9, 10).

We first assessed phagocytosis of *P. aeruginosa* PAO1 by THP-1 cells in the presence of mannose (Fig. 1A) and 2α-mannobiose (Fig. 1B), a disaccharide of mannose linked by a 1-2 glycosidic bond (31). Addition of either of these compounds resulted in significantly reduced uptake of *P. aeruginosa* by ∼50%. The addition of with N-acetylglucosamine at a relatively high concentration (Fig. 1C) and fucose (Fig. 1D) showed a statistically significant but modest (∼20%) reduction in *P. aeruginosa* phagocytosis. Finally, consistent with our previous observation (10), the internalization of *P. aeruginosa* was robustly and significantly diminished when the assay was supplemented with free galactose (Fig. 1E). Together, these data are consistent with the conclusion that *P. aeruginosa* PAO1 can bind to various mono- and di-saccharide components that comprise high-mannose and complex glycans (Fig. 1F).

**Figure 1.**
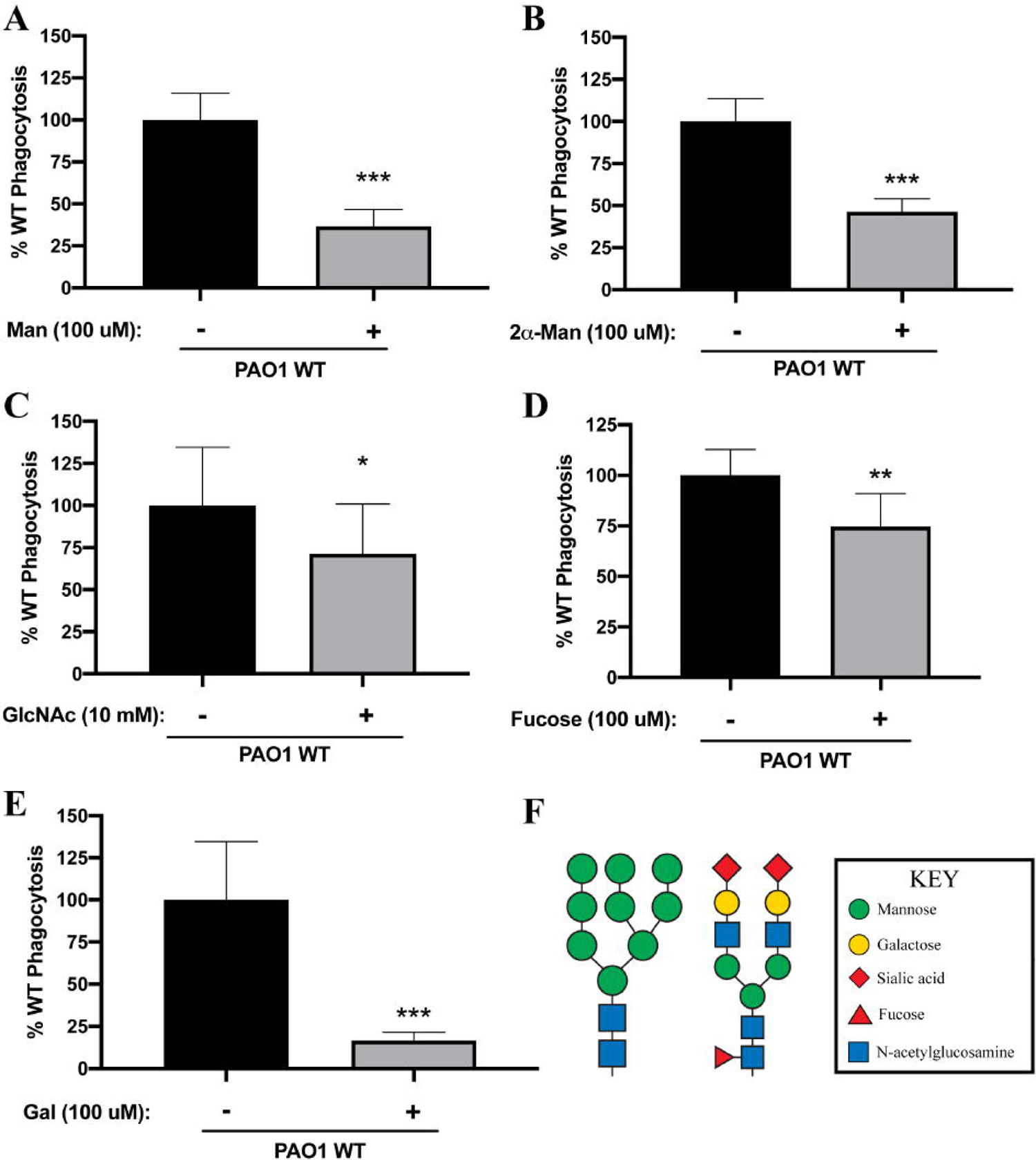
Competition with Mono- and Di-Saccharide Components of N-linked Glycans Inhibit *P. aeruginosa* PAO1 Phagocytosis. THP-1 cells were assayed for relative phagocytosis of *P. aeruginosa* PAO1 (MOI=10) in the absence or presence of the indicated, exogenously added sugars. Phagocytosis was normalized as a percentage of the mean of phagocytosis of *P. aeruginosa* by untreated THP-1 cells. (A-E) Mannose (Man; 100 µM), 2α-mannobiose (2α-Man; 100 µM), N-acetylglucosamine (GlcNAc; 10 mM), fucose (100 µM), and galactose (Gal; 100 µM) were assessed in the competition assays. Data in panels A-E were analyzed using an unpaired *t* test with Welch’s correction (A-E) and are representative of at least two independent biological experiments (n=8). ***, *P* ≤ 0.0005; **, *P* ≤ 0.005; *, P ≤ 0.05. (F) Schematic representations of N-linked structures: high-mannose type (left) and complex type (right). The component sugars comprising each glycan are indicated in the legend.

### An Assay to Quantitatively Profile Interactions of *P. aeruginosa* with Glycans

To identify additional candidate glycan ligands for *P. aeruginosa*, we utilized an otherwise wild-type, GFP-expressing strain of *P. aeruginosa* PAO1 and performed a bacterial binding assay with the commercially available Glycan Array 100 (RayBiotech). Each of the 100 glycan types is spotted in quadruplicate. An overnight culture of *P. aeruginosa* PAO1 carrying a GFP-expressing plasmid grown in lysogeny broth (LB), was sub-cultured and grown to mid-log phase in LB, then diluted 1:1000 to ∼1 x 10^6^ cells/ml into the manufacture’s minimal Sample Diluent Buffer. After a 3 hr incubation at room temperature with this bacterial suspension, each array was washed to remove non-binding cells, and fluorescence intensity values associated with the spotted glycans were quantified. These experiments were performed in triplicate, and additional details of the experiment and its analysis are presented in the Materials and Methods.

We observed a total of 28 glycans to which *P. aeruginosa* consistently and significantly bound (Figure 2A). These glycans are as follows: **1** β-Glc, **2** β-Gal, **3** α-Man, **4** α-Fuc, **5** α-Rha, **6** β-GlcNAc, **7** β-GalNAc, **8** Tobramycin, **9** Gal-β-1,3-GlcNAc-β, **10** Gal-α-1,3-Gal-β-1,3-GlcNAc-β, **11** Neu5Ac-α-2,3-Gal-β-1,3-GlcNAc-β, **20** GalNAc-β-1,3-Gal-β-1,4-Glc-β, **24** Neu5Gc-α-2,6-Gal-β-1,4-Glc-β, **25** Gal-β-1,4-(Fuc-α-1,3)-Glc-β, **27** GlcNAc-β-1,6-GlcNAc-β, **28** 4-P-GlcNAc-β-1,4-Man-β, **29** Glc-α-1,2-Gal-α-1,3-Glc-α, **37** Gal-β-1,4-GlcNAc-β-1,3-Gal-β-1,4-Glc-β-[LNnT], **38** GlcA-β-1,4-GlcNAc-α-1,4-GlcA-β, **41** GalNAc-β-1,4-GlcNAc-β, **48** Gal-α-1,2-Gal-α, **53** Gal-β-1,4-(6S)GlcNAc-β, **57** GalNAc-α-1,3-(Fuc-α-1,2)-Gal-β-[Blood A antigen trisaccharide], **59** Gal-α-1,3-(Fuc-α-1,2)-Gal-β-[Blood B antigen trisaccharide], **75** β-D-Rha-Sp, **76** Glc-α-1,4-Glc-β, **77** Glc-α-1,6-Glc-α-1,4-Glc-β, and **100** SGP (Sialylglycopeptide). A representative array after binding by *P. aeruginosa* is shown with both negative and positive fluorescence controls included is shown in Figure 2B. The quantification of binding of each of the 28 glycans is shown in Figure 2C, with the glycans ordered by signal intensity. A summary of the glycan array binding data is provided in Table 1.

**Fig. 2.**
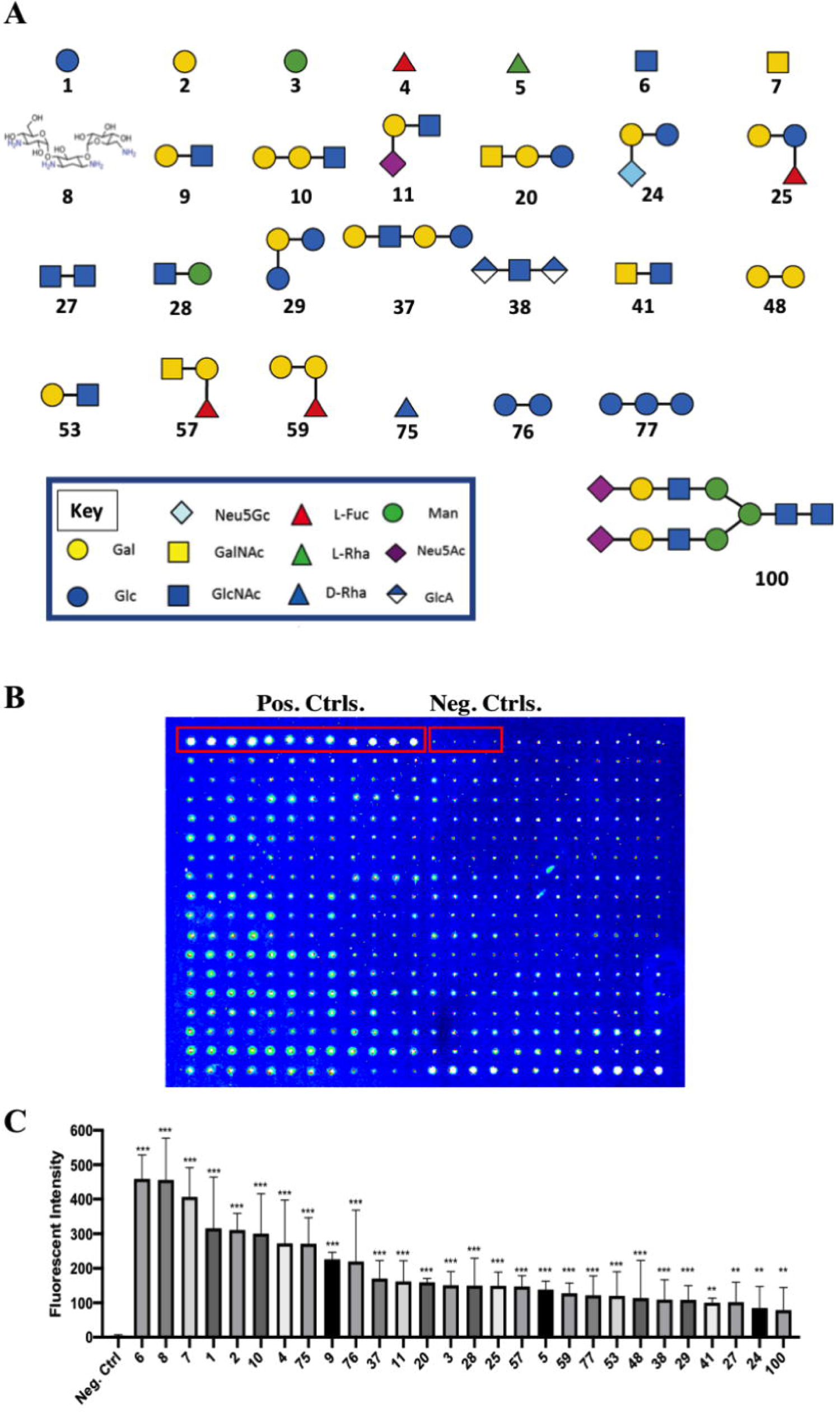
Analysis of Glycan Binding by *P. aeruginosa* PAO1 Using the Glycan Array 100. GFP-expressing *P. aeruginosa* PAO1 was quantitatively assessed for binding to known glycan structures printed on the Glycan Array 100. (A) Structures of the 28 compounds significantly bound by *P. aeruginosa* PAO1. The numbering corresponds to the text and to the Glycan Array 100. The key is shown at the bottom of the panel. The structures are modified from the figures at www.Raybiotech.com. (B) Representative example of one of the arrays tested for glycan binding by *P. aeruginosa*. The positive (a fluorescent protein) and negative (nothing spotted) controls are labeled and indicated by the red boxes. (C) Glycan array analysis revealed that *P. aeruginosa* PAO1 significantly bound 28 distinct glycans in this assay. The average fluorescence intensity associated with each glycan was compared to the negative control. The number on the X-axis corresponds to the numbering in the text and in panel A of this figure. Data were analyzed using a one-way ANOVA with Dunnett’s *post hoc* analyses and are derived from three independent experiments with quadruple spotting of each glycan on each array (n=12). ***, *P* ≤ 0.0005; **, *P* ≤ 0.005.

**Table 1.**
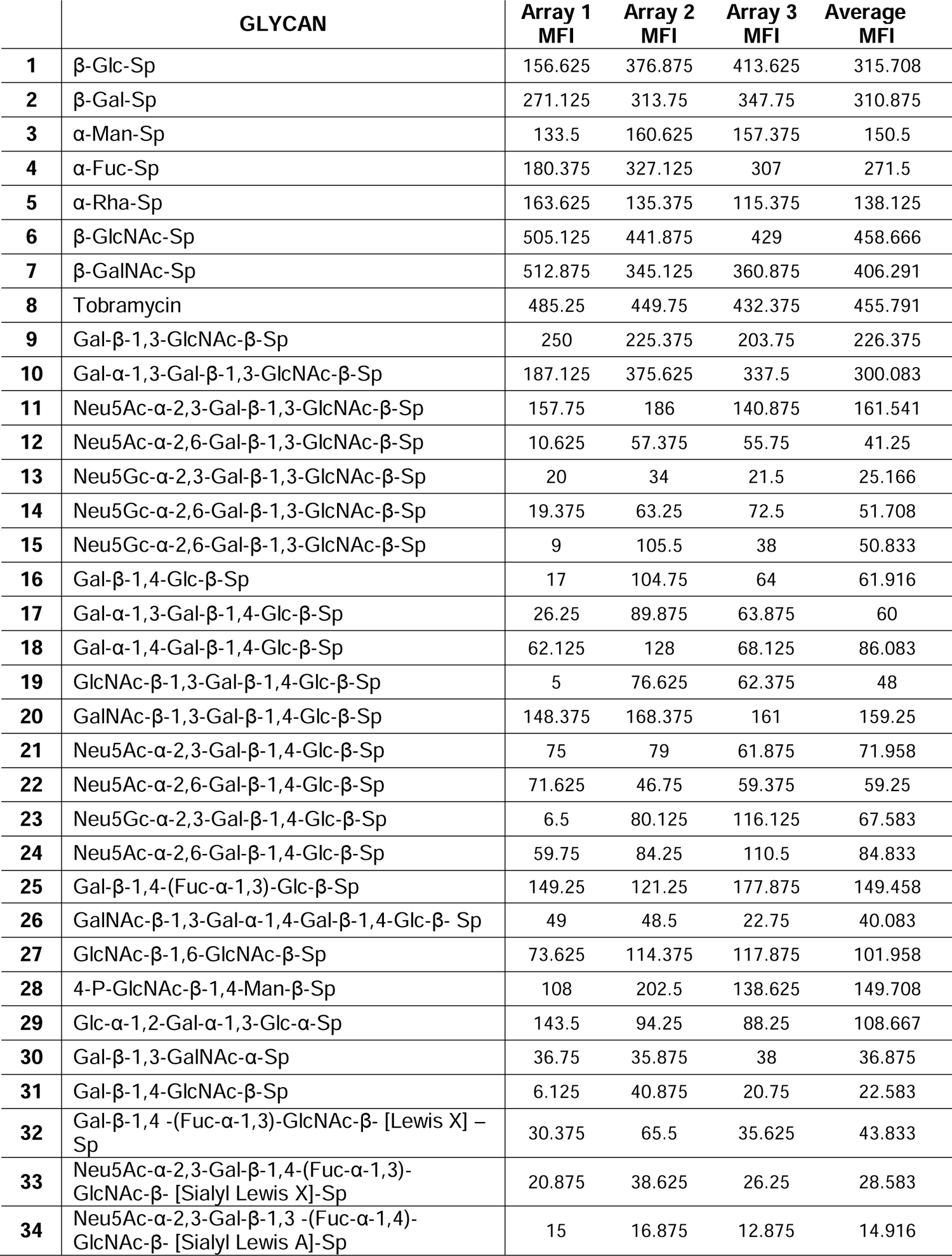
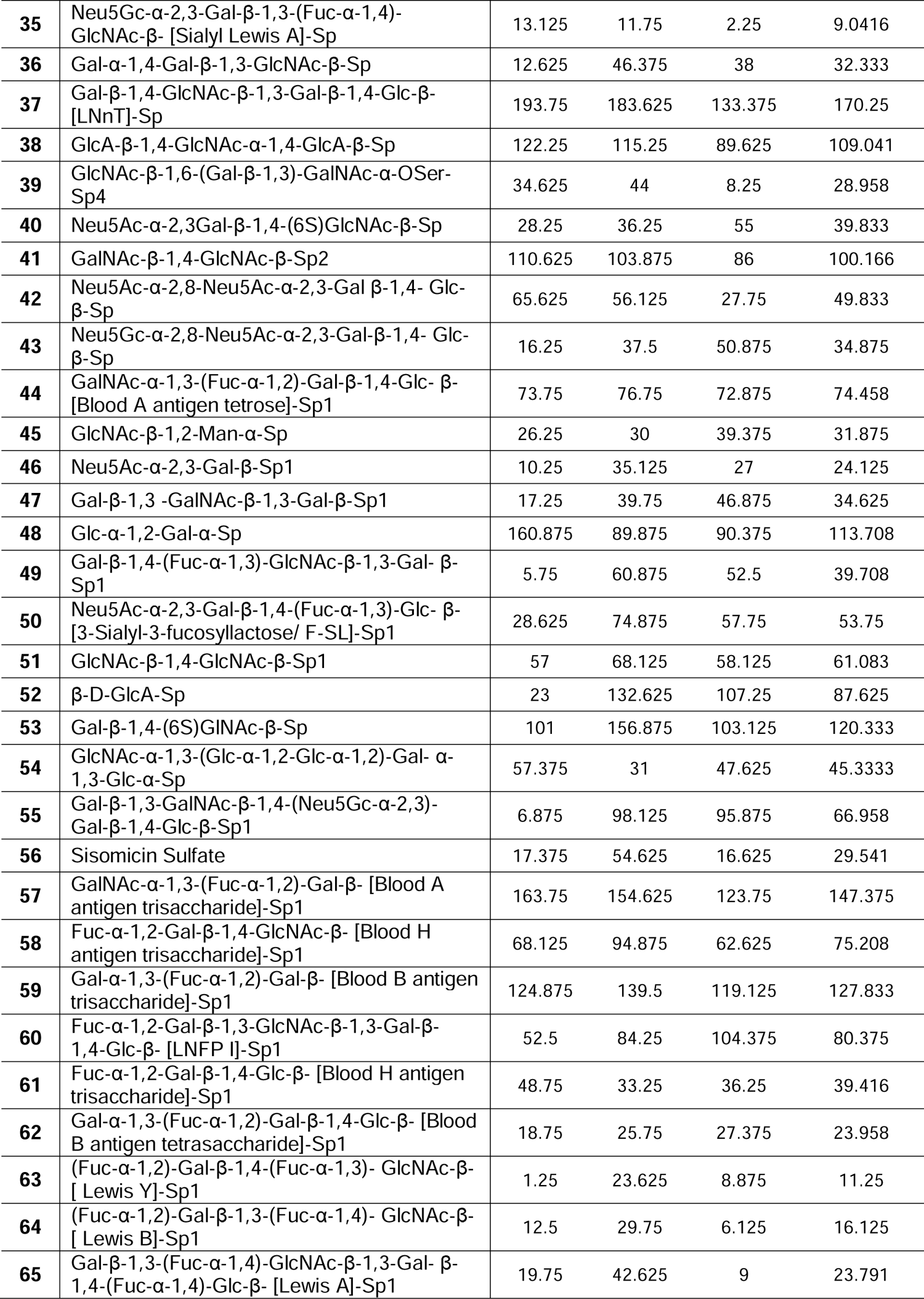
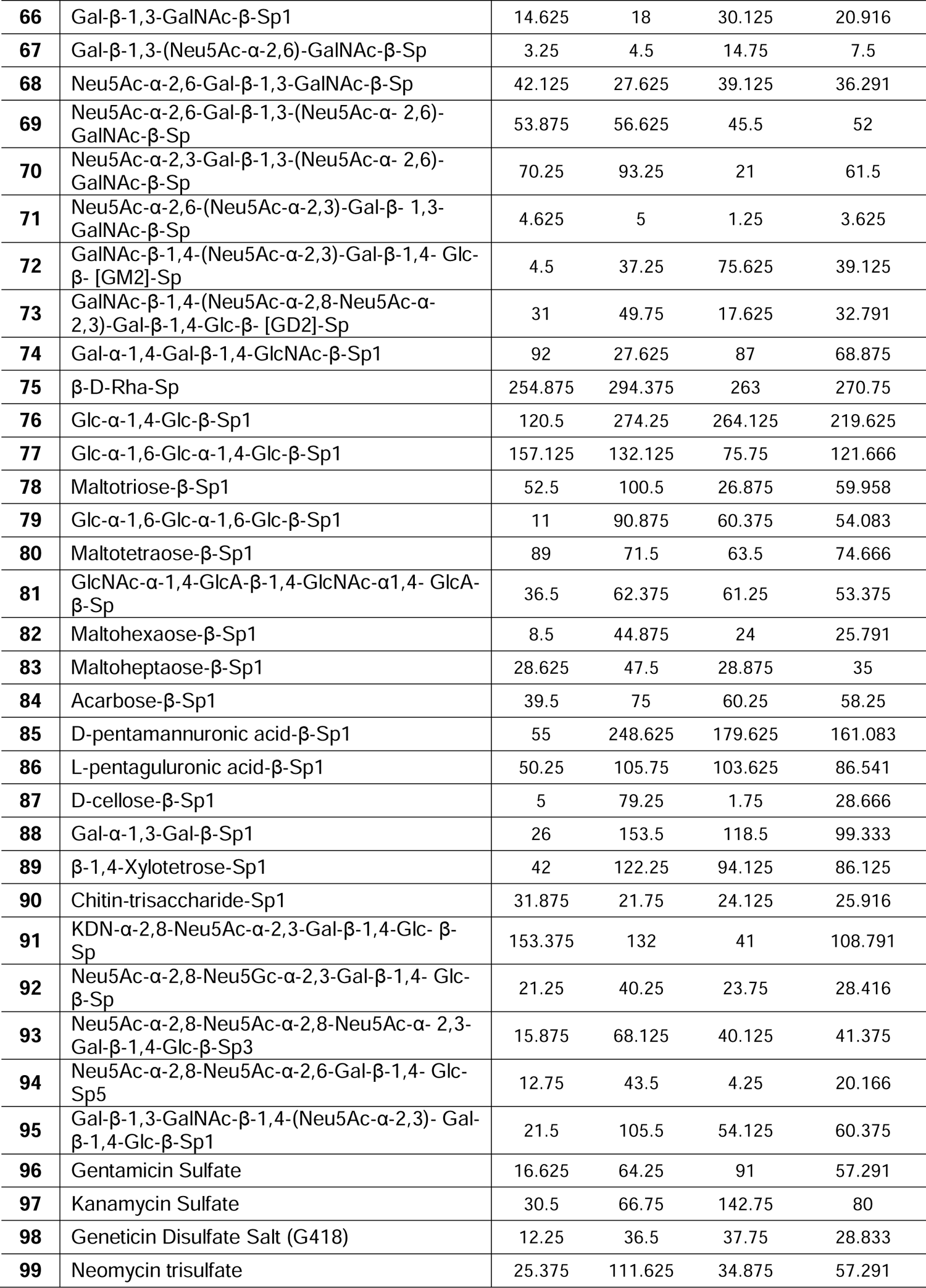
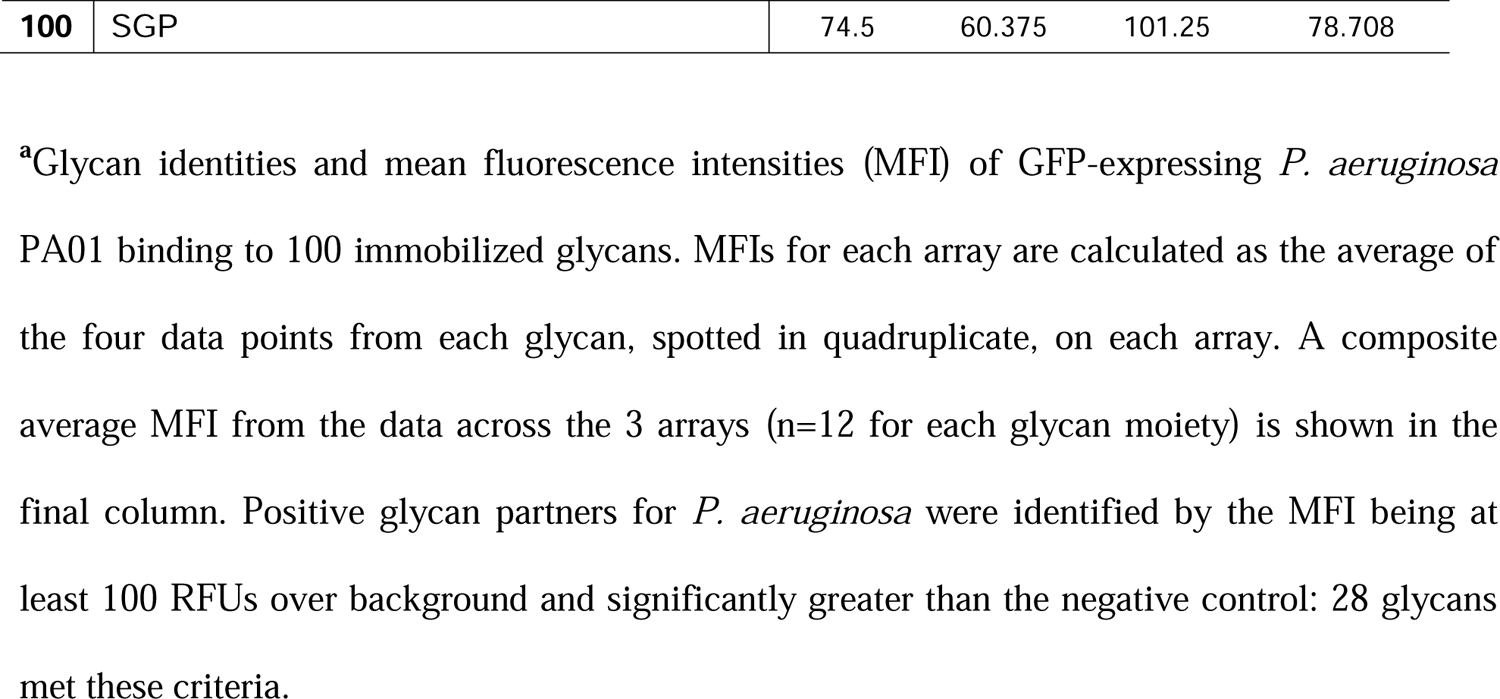
Glycan binding profile of *P. aeruginosa* PAO1.^a^

|  | GLYCAN | Array 1<br>MFI | Array 2<br>MFI | Array 3<br>MFI | Average<br>MFI |
| --- | --- | --- | --- | --- | --- |
| 1 | β-Glc-Sp | 156.625 | 376.875 | 413.625 | 315.708 |
| 2 | β-Gal-Sp | 271.125 | 313.75 | 347.75 | 310.875 |
| 3 | α-Man-Sp | 133.5 | 160.625 | 157.375 | 150.5 |
| 4 | α-Fuc-Sp | 180.375 | 327.125 | 307 | 271.5 |
| 5 | α-Rha-Sp | 163.625 | 135.375 | 115.375 | 138.125 |
| 6 | β-GlcNAc-Sp | 505.125 | 441.875 | 429 | 458.666 |
| 7 | β-GalNAc-Sp | 512.875 | 345.125 | 360.875 | 406.291 |
| 8 | Tobramycin | 485.25 | 449.75 | 432.375 | 455.791 |
| 9 | Gal-β-1,3-GlcNAc-β-Sp | 250 | 225.375 | 203.75 | 226.375 |
| 10 | Gal-α-1,3-Gal-β-1,3-GlcNAc-β-Sp | 187.125 | 375.625 | 337.5 | 300.083 |
| 11 | Neu5Ac-α-2,3-Gal-β-1,3-GlcNAc-β-Sp | 157.75 | 186 | 140.875 | 161.541 |
| 12 | Neu5Ac-α-2,6-Gal-β-1,3-GlcNAc-β-Sp | 10.625 | 57.375 | 55.75 | 41.25 |
| 13 | Neu5Gc-α-2,3-Gal-β-1,3-GlcNAc-β-Sp | 20 | 34 | 21.5 | 25.166 |
| 14 | Neu5Gc-α-2,6-Gal-β-1,3-GlcNAc-β-Sp | 19.375 | 63.25 | 72.5 | 51.708 |
| 15 | Neu5Gc-α-2,6-Gal-β-1,3-GlcNAc-β-Sp | 9 | 105.5 | 38 | 50.833 |
| 16 | Gal-β-1,4-Glc-β-Sp | 17 | 104.75 | 64 | 61.916 |
| 17 | Gal-α-1,3-Gal-β-1,4-Glc-β-Sp | 26.25 | 89.875 | 63.875 | 60 |
| 18 | Gal-α-1,4-Gal-β-1,4-Glc-β-Sp | 62.125 | 128 | 68.125 | 86.083 |
| 19 | GlcNAc-β-1,3-Gal-β-1,4-Glc-β-Sp | 5 | 76.625 | 62.375 | 48 |
| 20 | GalNAc-β-1,3-Gal-β-1,4-Glc-β-Sp | 148.375 | 168.375 | 161 | 159.25 |
| 21 | Neu5Ac-α-2,3-Gal-β-1,4-Glc-β-Sp | 75 | 79 | 61.875 | 71.958 |
| 22 | Neu5Ac-α-2,6-Gal-β-1,4-Glc-β-Sp | 71.625 | 46.75 | 59.375 | 59.25 |
| 23 | Neu5Gc-α-2,3-Gal-β-1,4-Glc-β-Sp | 6.5 | 80.125 | 116.125 | 67.583 |
| 24 | Neu5Ac-α-2,6-Gal-β-1,4-Glc-β-Sp | 59.75 | 84.25 | 110.5 | 84.833 |
| 25 | Gal-β-1,4-(Fuc-α-1,3)-Glc-β-Sp | 149.25 | 121.25 | 177.875 | 149.458 |
| 26 | GalNAc-β-1,3-Gal-α-1,4-Gal-β-1,4-Glc-β-Sp | 49 | 48.5 | 22.75 | 40.083 |
| 27 | GlcNAc-β-1,6-GlcNAc-β-Sp | 73.625 | 114.375 | 117.875 | 101.958 |
| 28 | 4-P-GlcNAc-β-1,4-Man-β-Sp | 108 | 202.5 | 138.625 | 149.708 |
| 29 | Glc-α-1,2-Gal-α-1,3-Glc-α-Sp | 143.5 | 94.25 | 88.25 | 108.667 |
| 30 | Gal-β-1,3-GalNAc-α-Sp | 36.75 | 35.875 | 38 | 36.875 |
| 31 | Gal-β-1,4-GlcNAc-β-Sp | 6.125 | 40.875 | 20.75 | 22.583 |
| 32 | Gal-β-1,4-(Fuc-α-1,3)-GlcNAc-β-[Lewis X]-Sp | 30.375 | 65.5 | 35.625 | 43.833 |
| 33 | Neu5Ac-α-2,3-Gal-β-1,4-(Fuc-α-1,3)-GlcNAc-β-[Sialyl Lewis X]-Sp | 20.875 | 38.625 | 26.25 | 28.583 |
| 34 | Neu5Ac-α-2,3-Gal-β-1,3-(Fuc-α-1,4)-GlcNAc-β-[Sialyl Lewis A]-Sp | 15 | 16.875 | 12.875 | 14.916 |
| 35 | Neu5Gc-α-2,3-Gal-β-1,3-(Fuc-α-1,4)-GlcNAc-β- [Sialyl Lewis A]-Sp | 13.125 | 11.75 | 2.25 | 9.0416 |
| 36 | Gal-α-1,4-Gal-β-1,3-GlcNAc-β-Sp | 12.625 | 46.375 | 38 | 32.333 |
| 37 | Gal-β-1,4-GlcNAc-β-1,3-Gal-β-1,4-Glc-β-[LNnT]-Sp | 193.75 | 183.625 | 133.375 | 170.25 |
| 38 | GlcA-β-1,4-GlcNAc-α-1,4-GlcA-β-Sp | 122.25 | 115.25 | 89.625 | 109.041 |
| 39 | GlcNAc-β-1,6-(Gal-β-1,3)-GalNAc-α-OSer-Sp4 | 34.625 | 44 | 8.25 | 28.958 |
| 40 | Neu5Ac-α-2,3Gal-β-1,4-(6S)GlcNAc-β-Sp | 28.25 | 36.25 | 55 | 39.833 |
| 41 | GalNAc-β-1,4-GlcNAc-β-Sp2 | 110.625 | 103.875 | 86 | 100.166 |
| 42 | Neu5Ac-α-2,8-Neu5Ac-α-2,3-Gal β-1,4- Glc-β-Sp | 65.625 | 56.125 | 27.75 | 49.833 |
| 43 | Neu5Gc-α-2,8-Neu5Ac-α-2,3-Gal-β-1,4- Glc-β-Sp | 16.25 | 37.5 | 50.875 | 34.875 |
| 44 | GalNAc-α-1,3-(Fuc-α-1,2)-Gal-β-1,4-Glc- β-[Blood A antigen tetrose]-Sp1 | 73.75 | 76.75 | 72.875 | 74.458 |
| 45 | GlcNAc-β-1,2-Man-α-Sp | 26.25 | 30 | 39.375 | 31.875 |
| 46 | Neu5Ac-α-2,3-Gal-β-Sp1 | 10.25 | 35.125 | 27 | 24.125 |
| 47 | Gal-β-1,3 -GalNAc-β-1,3-Gal-β-Sp1 | 17.25 | 39.75 | 46.875 | 34.625 |
| 48 | Glc-α-1,2-Gal-α-Sp | 160.875 | 89.875 | 90.375 | 113.708 |
| 49 | Gal-β-1,4-(Fuc-α-1,3)-GlcNAc-β-1,3-Gal- β-Sp1 | 5.75 | 60.875 | 52.5 | 39.708 |
| 50 | Neu5Ac-α-2,3-Gal-β-1,4-(Fuc-α-1,3)-Glc- β-[3-Sialyl-3-fucosyllactose/ F-SL]-Sp1 | 28.625 | 74.875 | 57.75 | 53.75 |
| 51 | GlcNAc-β-1,4-GlcNAc-β-Sp1 | 57 | 68.125 | 58.125 | 61.083 |
| 52 | β-D-GlcA-Sp | 23 | 132.625 | 107.25 | 87.625 |
| 53 | Gal-β-1,4-(6S)GINAc-β-Sp | 101 | 156.875 | 103.125 | 120.333 |
| 54 | GlcNAc-α-1,3-(Glc-α-1,2-Glc-α-1,2)-Gal- α-1,3-Glc-α-Sp | 57.375 | 31 | 47.625 | 45.3333 |
| 55 | Gal-β-1,3-GalNAc-β-1,4-(Neu5Gc-α-2,3)-Gal-β-1,4-Glc-β-Sp1 | 6.875 | 98.125 | 95.875 | 66.958 |
| 56 | Sisomicin Sulfate | 17.375 | 54.625 | 16.625 | 29.541 |
| 57 | GalNAc-α-1,3-(Fuc-α-1,2)-Gal-β- [Blood A antigen trisaccharide]-Sp1 | 163.75 | 154.625 | 123.75 | 147.375 |
| 58 | Fuc-α-1,2-Gal-β-1,4-GlcNAc-β- [Blood H antigen trisaccharide]-Sp1 | 68.125 | 94.875 | 62.625 | 75.208 |
| 59 | Gal-α-1,3-(Fuc-α-1,2)-Gal-β- [Blood B antigen trisaccharide]-Sp1 | 124.875 | 139.5 | 119.125 | 127.833 |
| 60 | Fuc-α-1,2-Gal-β-1,3-GlcNAc-β-1,3-Gal-β-1,4-Glc-β- [LNFP I]-Sp1 | 52.5 | 84.25 | 104.375 | 80.375 |
| 61 | Fuc-α-1,2-Gal-β-1,4-Glc-β- [Blood H antigen trisaccharide]-Sp1 | 48.75 | 33.25 | 36.25 | 39.416 |
| 62 | Gal-α-1,3-(Fuc-α-1,2)-Gal-β-1,4-Glc-β- [Blood B antigen tetrasaccharide]-Sp1 | 18.75 | 25.75 | 27.375 | 23.958 |
| 63 | (Fuc-α-1,2)-Gal-β-1,4-(Fuc-α-1,3)- GlcNAc-β-[Lewis Y]-Sp1 | 1.25 | 23.625 | 8.875 | 11.25 |
| 64 | (Fuc-α-1,2)-Gal-β-1,3-(Fuc-α-1,4)- GlcNAc-β-[Lewis B]-Sp1 | 12.5 | 29.75 | 6.125 | 16.125 |
| 65 | Gal-β-1,3-(Fuc-α-1,4)-GlcNAc-β-1,3-Gal- β-1,4-(Fuc-α-1,4)-Glc-β- [Lewis A]-Sp1 | 19.75 | 42.625 | 9 | 23.791 |
| 66 | Gal-β-1,3-GalNAc-β-Sp1 | 14.625 | 18 | 30.125 | 20.916 |
| 67 | Gal-β-1,3-(Neu5Ac-α-2,6)-GalNAc-β-Sp | 3.25 | 4.5 | 14.75 | 7.5 |
| 68 | Neu5Ac-α-2,6-Gal-β-1,3-GalNAc-β-Sp | 42.125 | 27.625 | 39.125 | 36.291 |
| 69 | Neu5Ac-α-2,6-Gal-β-1,3-(Neu5Ac-α-2,6)-GalNAc-β-Sp | 53.875 | 56.625 | 45.5 | 52 |
| 70 | Neu5Ac-α-2,3-Gal-β-1,3-(Neu5Ac-α-2,6)-GalNAc-β-Sp | 70.25 | 93.25 | 21 | 61.5 |
| 71 | Neu5Ac-α-2,6-(Neu5Ac-α-2,3)-Gal-β-1,3-GalNAc-β-Sp | 4.625 | 5 | 1.25 | 3.625 |
| 72 | GalNAc-β-1,4-(Neu5Ac-α-2,3)-Gal-β-1,4-Glc-β-[GM2]-Sp | 4.5 | 37.25 | 75.625 | 39.125 |
| 73 | GalNAc-β-1,4-(Neu5Ac-α-2,8-Neu5Ac-α-2,3)-Gal-β-1,4-Glc-β-[GD2]-Sp | 31 | 49.75 | 17.625 | 32.791 |
| 74 | Gal-α-1,4-Gal-β-1,4-GlcNAc-β-Sp1 | 92 | 27.625 | 87 | 68.875 |
| 75 | β-D-Rha-Sp | 254.875 | 294.375 | 263 | 270.75 |
| 76 | Glc-α-1,4-Glc-β-Sp1 | 120.5 | 274.25 | 264.125 | 219.625 |
| 77 | Glc-α-1,6-Glc-α-1,4-Glc-β-Sp1 | 157.125 | 132.125 | 75.75 | 121.666 |
| 78 | Maltotriose-β-Sp1 | 52.5 | 100.5 | 26.875 | 59.958 |
| 79 | Glc-α-1,6-Glc-α-1,6-Glc-β-Sp1 | 11 | 90.875 | 60.375 | 54.083 |
| 80 | Maltotetraose-β-Sp1 | 89 | 71.5 | 63.5 | 74.666 |
| 81 | GlcNAc-α-1,4-GlcA-β-1,4-GlcNAc-α-1,4-GlcA-β-Sp | 36.5 | 62.375 | 61.25 | 53.375 |
| 82 | Maltohexaose-β-Sp1 | 8.5 | 44.875 | 24 | 25.791 |
| 83 | Maltoheptaose-β-Sp1 | 28.625 | 47.5 | 28.875 | 35 |
| 84 | Acarbose-β-Sp1 | 39.5 | 75 | 60.25 | 58.25 |
| 85 | D-pentamannuronic acid-β-Sp1 | 55 | 248.625 | 179.625 | 161.083 |
| 86 | L-pentaguluronic acid-β-Sp1 | 50.25 | 105.75 | 103.625 | 86.541 |
| 87 | D-cellose-β-Sp1 | 5 | 79.25 | 1.75 | 28.666 |
| 88 | Gal-α-1,3-Gal-β-Sp1 | 26 | 153.5 | 118.5 | 99.333 |
| 89 | β-1,4-Xylotetrose-Sp1 | 42 | 122.25 | 94.125 | 86.125 |
| 90 | Chitin-trisaccharide-Sp1 | 31.875 | 21.75 | 24.125 | 25.916 |
| 91 | KDN-α-2,8-Neu5Ac-α-2,3-Gal-β-1,4-Glc-β-Sp | 153.375 | 132 | 41 | 108.791 |
| 92 | Neu5Ac-α-2,8-Neu5Gc-α-2,3-Gal-β-1,4-Glc-β-Sp | 21.25 | 40.25 | 23.75 | 28.416 |
| 93 | Neu5Ac-α-2,8-Neu5Ac-α-2,8-Neu5Ac-α-2,3-Gal-β-1,4-Glc-β-Sp3 | 15.875 | 68.125 | 40.125 | 41.375 |
| 94 | Neu5Ac-α-2,8-Neu5Ac-α-2,6-Gal-β-1,4-Glc-Sp5 | 12.75 | 43.5 | 4.25 | 20.166 |
| 95 | Gal-β-1,3-GalNAc-β-1,4-(Neu5Ac-α-2,3)-Gal-β-1,4-Glc-β-Sp1 | 21.5 | 105.5 | 54.125 | 60.375 |
| 96 | Gentamicin Sulfate | 16.625 | 64.25 | 91 | 57.291 |
| 97 | Kanamycin Sulfate | 30.5 | 66.75 | 142.75 | 80 |
| 98 | Geneticin Disulfate Salt (G418) | 12.25 | 36.5 | 37.75 | 28.833 |
| 99 | Neomycin trisulfate | 25.375 | 111.625 | 34.875 | 57.291 |
| **100** | SGP | 74.5 | 60.375 | 101.25 | 78.708 |

<sup>a</sup>Glycan identities and mean fluorescence intensities (MFI) of GFP-expressing *P. aeruginosa* PA01 binding to 100 immobilized glycans. MFIs for each array are calculated as the average of the four data points from each glycan, spotted in quadruplicate, on each array. A composite average MFI from the data across the 3 arrays (n=12 for each glycan moiety) is shown in the final column. Positive glycan partners for *P. aeruginosa* were identified by the MFI being at least 100 RFUs over background and significantly greater than the negative control: 28 glycans met these criteria.

### *Pseudomonas aeruginosa* Binds to Monosaccharides and an Aminoglycoside

We further examined the binding results for each of the glycan classes spotted on the Glycan Array 100 to identify binding signatures. The glycans on the array can be divided into eight groups: (A) monosaccharides, (B) aminoglycosides, (C) disaccharides, (D) blood groups, Lewis antigens, and fucosylated oligosaccharides, (E) globo-series glycolipids, milk oligosaccharides, and GAGs, (F) natural oligosaccharides, (G) gangliosides and sialylated oligosaccharides, and (H) α-Gal & N-glycans. Notably, a majority of positive glycan signatures identified were to monosaccharides and disaccharides: 15 of the 28 significantly bound glycans fell into this category. This analysis also revealed that 14 out of the 28 glycans identified had at least one galactose moiety. Furthermore, 11 of the 28 glycans to which *P. aeruginosa* bound contained at least one GlcNAc moiety. We describe the binding of *P. aeruginosa* PAO1 to each specific glycan class below.

The significant attachment of *P. aeruginosa* to monosaccharides included: 1, glucose; 2, galactose; 3, mannose; 4, fucose; 5, rhamnose; 6, GlcNAc; 7, GalNAc (Fig. 3A). With the exception of glucose and rhamnose, these glycans are all components of N-linked glycan structures and are capable of competing for bacterial binding sites on phagocytes (Fig. 1). Furthermore, *P. aeruginosa* significantly bound tobramycin (Fig. 3B), an antibiotic used for the treatment of this microbe.

**Fig. 3.**
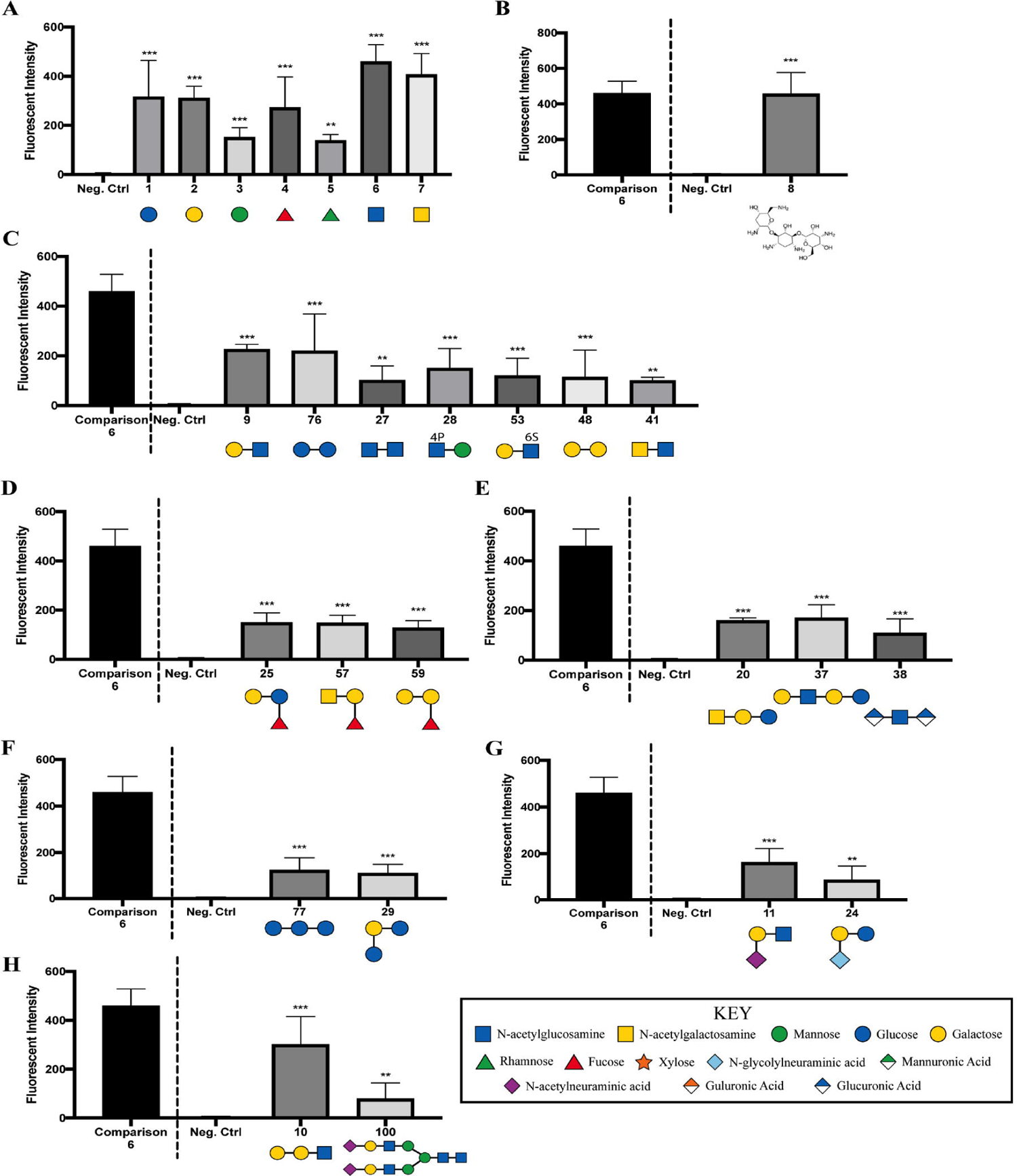
Glycan Groups that Exhibit Significant Binding by *P. aeruginosa* PAO1. Glycan classes significantly bound by *P. aeruginosa* PAO1 as measured using the Glycan Array 100: (A) monosaccharides, (B) an aminoglycoside, (C) disaccharides, (D) blood groups, Lewis antigens, and fucosylated oligosaccharides, (E) globo-series glycolipids, milk oligosaccharides, and GAGs, (F) natural oligosaccharides, (G) gangliosides and sialylated oligosaccharides, and (H) α-Gal and N-glycans. Data were normalized to binding of GFP-expressing *P. aeruginosa* PAO1 compared to the negative control and dotted line separates glycan **6** (GlcNAc), which serves as a point of comparison with the other tested glycans. Data were analyzed using a one-way ANOVA with Dunnett’s *post hoc* analyses (A, C-H) or an unpaired *t* test with Welch’s correction (B) and are representative of three independent experiments with each glycan spotted in quadruplicate (n=12). ***, *P* ≤ 0.0005; **, *P* ≤ 0.005.

*P. aeruginosa* showed the most robust binding to Gal-β-1,3-GlcNAc (Glycan **9**) and Glc-α-1,4-Glc-β (Glycan **76**) in comparison to other disaccharides (Fig. 3C). Gal-β-1,3-GlcNAc has been described in complex and hybrid types of N-linked glycosylation (29). *P. aeruginosa* bound to the diamino-saccharide GlcNAc-β-1,6-GlcNAc (Glycan **27**) to a lesser extent than to the mono-aminosaccharide GlcNAc (Glycan **6**; Fig. 3A,C).

In addition to N-linked associated glycans, *P. aeruginosa* PAO1 also binds to selected: blood groups, Lewis antigens, and fucosylated oligosaccharides (Fig. 3D), globo-series glycolipids, milk oligosaccharides, and GAGs (Fig. 3E), natural oligosaccharides (Fig. 3F), gangliosides and sialylated oligosaccharides (Fig. 3G), and α-Gal and N-glycans (Fig. 3H).

## DISCUSSION

*P. aeruginosa* can bind to glycans in a variety of contexts. Previous studies have shown that glycan moieties contribute to *P. aeruginosa* binding of airway epithelium (18, 19), and furthermore, bacterial surface adhesins can mediate binding to glycan ligands on host cells (32–35) and on other bacterial species (13, 15, 36). Additionally, recent studies revealed that the inhibition of N-linked glycan synthesis on host innate immune cells by tunicamycin decreased bacterial binding and, consequently, phagocytosis of *P. aeruginosa* and IL-1β elicitation from the host cells (10). However, studies to date have interrogated a limited number of glycans that do not encompass all of the possible complexity of glycan structures (10, 11, 36). In this work, we combine cellular assays based on knowledge gained from previous studies with a screen to identify additional candidate glycans bound by *P. aeruginosa*.

Since *P. aeruginosa* preferentially binds to N-linked glycans (10), as illustrated in Figure 1F, initial efforts were focused on the identification of components of these glycans that may confer binding. Competition studies for bacterial binding and uptake revealed that specific monosaccharide components of N-linked glycans, including mannose and galactose, are effective at competing bacterial association with THP-1 monocytic cells while, in comparison, other monosaccharides, including fucose, at equimolar concentrations, are much less effective.

To further identify the breadth of glycans bound by *P. aeruginosa*, we performed quantitative binding assay with GFP-labeled P. aeruginosa using the Glycan 100 Array. Interestingly, we found that the majority of bound glycans have at one or more galactose moieties, which aligns with the competition studies presented here (Figure 1). We note that galactose was found in the following glycans identified in the array assay: Gal-β-1,4-(Fuc-α-1,3)-Glc (Glycan **25**), Blood A antigen (Glycan **57**), Blood B antigen (Glycan **59**), GalNAc-β-1,3-Gal-β-1,4-Glc (Glycan **20**), Gal-β-1,4-GlcNAc-β-1,3-Gal-β-1,4-Glc (Glycan **37**), Glc-α-1,2-Gal-α-1,3-Glc (Glycan **29**), Neu5Ac-α-2,3-Gal-β-1,3-GlcNAc (Glycan **11**), Neu5Gc-α-2,6-Gal-β-1,4-Glc (Glycan **24**), Gal-α-1,3-Gal-β-1,3-GlcNAc (Glycan **10**), and SGP (Glycan **100**). Similarly, mannose, identified in the competition assays was also bound by *P. aeruginosa* in the array study, and GlcNac was shown to mediate robust binding by *P. aeruginosa* despite only a modest effect on phagocytosis in the competition assays (Fig. 1). Overall, from the eight specified classes of glycans outlined above, monosaccharides, disaccharides, and an aminoglycoside were featured among the compounds bound by *P. aeruginosa*. Finally, we note that one limitation of this assay is that we cannot completely rule out some bacterial growth during the three hour incubation from carry-over of nutrients during the dilution of the culture 1:1000 into the Sample Diluent Buffer. Any such growth would simply enhance the relative binding signal, as growing bacteria that were not adhered to the array would be removed during the washing step.

How do our findings fit within the context of previous studies? A previous study by Ramphal and colleagues showed that *P. aeruginosa* binds two different Gal-GlcNAc disaccharides (25), consistent with our findings here (Fig. 2, compounds **9** and **53**). Similarly, this microbe binds to the GalNAcβ1-4Gal disaccharide (37), also consistent with our findings. Previous work also indicated that *P. aeruginosa* can bind *N*-acetylglucosamine and sialic acids (21), lipid-linked lactose and lactose-derivatives (24), and sialyl-Lewis x conjugates (22). We confirmed many of these findings here, supporting previous studies that showed the ability of *P. aeruginosa* to bind to specific moieties of glycolipids and mucins (20, 25), and serving as a validation of our studies. *P. aeruginosa* has also been shown to bind to chitin (27), but we did not address binding to this sugar polymer.

One particularly intriguing observation was that *P. aeruginosa* binds to tobramycin (Glycan **8**). Tobramycin is a broad-spectrum antibiotic that is administered via intravenous or intramuscular injection, and is also utilized heavily by oral inhalation for treatment the of CF-associated chronic infections (38). Tobramycin inhibits protein synthesis by binding to the 16S ribosomal RNA of the bacterial 30S ribosome, which leads to mistranslation and subsequent cell membrane damage (39). This study suggests a novel mechanism of interaction between *P. aeruginosa* and this antibiotic, perhaps explaining its potency in killing this microbe (40). Additional studies will be needed to delineate how the observed binding may contribute to tobramycin’s antibiotic activity.

## Materials and Methods

### Bacteria

*P. aeruginosa* strain PAO1 carry GFP-expressing plasmid pSMC21 (41) and provided by Dr. D. Hogan was used here and has been used in previous studies (4, 5, 42, 43). Bacteria were cultured overnight in LB at 37°C, subsequently sub-cultured to achieve log-phase growth for 2 h in LB, and bacterial growth determined by optical density at 600 nm.

### Cell culture

THP-1 human monocytic cells were provided by P. Guyre (Geisel School of Medicine at Dartmouth, Lebanon, NH). Using a modification of a previously published protocol (44), cells were cultured and maintained in RPMI 1640 medium (HyClone) supplemented with 10% fetal bovine serum (FBS; HyClone), 5% Pen/Strep, 5% L-glutamine and 1mM sodium pyruvate until harvest. The phagocytic cells were not activated.

### Gentamicin protection assay

Phagocytosis of live bacteria was performed and quantitated as previously described (4, 5, 9, 10). Briefly, overnight cultures of *P. aeruginosa* were washed and resuspended in serum-free Hanks balanced salt solution (HBSS; Corning) and added to the THP-1 cells at a multiplicity of infection (MOI) of 10. Where indicated, phagocytic cells were pretreated with the specified glycan for competition studies, as previously described (10). After the 45-min co-incubation of bacteria with the glycan or control, 100 µg/ml gentamicin was added to the assay for 20 min at 37°C. The cultures were washed and subsequently lysed in 500 µl 0.1% Triton X-100 solution in 1X PBS. Lysates were plated on LB plates and incubated overnight at 37°C to determine bacterial viable counts. The next day, recovered colony forming units (CFU) on LB plates were enumerated and represented as the percentage of the mean of WT bacteria phagocytosed or fold increase in phagocytosis, as indicated in the figure legends, to quantitatively compare relative phagocytosis levels.

### Glycan 100 Array analysis

The commercially-available Glycan 100 Array (RayBiotech) was used to assess carbohydrate binding preferences to 100 described glycan structures by *P. aeruginosa*. To assess glycan binding by *P. aeruginosa*, live GFP-expressing *P. aeruginosa* PAO1 cells were subjected to glycan array analysis (*n* = 3). After growth overnight in LB medium, the culture of *P. aeruginosa* PAO1 expressing GFP, sub-cultured into fresh LB for 2 ∼hrs, then diluted 1:1000 into the manufacturer’s Sample Diluent Buffer for a final bacterial cell count of ∼1 x 10^6^ CFU/ml. This suspension was applied to the array and incubated at room temperature for 3 hours as per the manufacture’s directions. The positive control is a fluorescent protein and the negative control has nothing spotted on the array. Scanning and analysis of the slide was performed using the DNA Microarray Scanner (Agilent Technologies).

The list of the glycans on the Glycan 100 Array is provided in Table 1. Glycan identities and mean fluorescence intensity (MFI) of GFP-expressing *P. aeruginosa* PAO1 binding to the glycans was determined. MFI for each glycan is calculated as the average of the four data points from each glycan (i.e., each glycan is spotted in quadruplicate on each array) from across the 3 arrays (thus n=12 for each glycan moiety). This average MFI is shown in the final column of Table 1. Positive glycan binding partners for *P. aeruginosa* were set using an MFI at >100 RFUs over background and significantly greater than the negative control: 28 glycans met these criteria.

### Statistical analyses

Means ± standard deviations (SD) derived from multiple independent experiments with technical replicates are shown for each graph. Sample sizes for each experiment are noted in the figure legends. As indicated in the legends, unpaired Student’s *t* test with Welch’s correction or one-way analysis of variance (ANOVA) with Tukey’s *post hoc* analyses were performed using Prism, version 7.02, to determine statistical significance of the data. Statistical significance is represented in the figures by asterisks.

## DATA AVAILABILITY STATEMENT

The one large data set, binding of *P. aeruginosa* to the glycan array, is available in the manuscript in Table 1.

## ACKNOWLEDGMENTS

We thank Drs. Deborah Hogan, William Rigby, Margie Ackerman, Robert Cramer, Jim Bliska, Lynn Theprungsirikul, Nicole Loeven, and Tammara Wood for reagents, assistance with array scanning, and discussions. This work was supported by grants from the National Institutes of Health (R21 AI137656, R21 AI121820, R03 AI135358 to B.B., T32 AI007363 to H.S., to R37 AI83256 G.O.). The funders had no role in study design, data collection and interpretation, or the decision to submit the work for publication.

